# Extreme drought triggers transition to an alternative soil microbial state

**DOI:** 10.1101/2021.12.10.472086

**Authors:** Irene Cordero, Ainara Leizeaga, Lettice C. Hicks, Johannes Rousk, Richard D. Bardgett

## Abstract

Soil microbial communities play a pivotal role in regulating ecosystem functioning^1^ but they are increasingly threatened by human-driven perturbations, including climate extremes, which are predicted to increase in frequency and intensity with climate change^2^. It has been demonstrated that soil microbial communities are sensitive to climate extremes, such as drought^3,4^, and that effects can be long-lasting^5,6^. However, considerable uncertainties remain concerning the response of soil microbial communities to increases in the intensity and frequency of climate extremes, and their potential to trigger transitions to alternative, and potentially deleterious, taxonomic and functional states^7^. Here we demonstrate that extreme, frequent drought induces a shift to an alternative soil microbial state characterised by strongly altered bacterial and fungal community structure of reduced complexity and functionality. Moreover, we found that this drought-induced alternative microbial state persisted after returning soil to its previous moisture status. However, bacterial communities were able to adapt by increasing their growth capacity, despite being of reduced diversity. Abrupt transitions to alternative states are well documented in aquatic and terrestrial plant communities in response to human-induced perturbations, including climate extremes^8,9^. Our results provide experimental evidence that such transitions also occur in soil microbial communities in response to extreme drought with potentially deleterious consequences for soil health.

## MAIN

Natural ecosystems are constantly exposed to natural fluctuations in environmental conditions and under such conditions they retain a stable equilibrium state, or quasi-stable state, characterized by minor fluctuations in community composition and function^10^. However, human-induced perturbations, such as those related to climate change, can destabilise this dynamic equilibrium and potentially trigger a cascade of events that may lead to an abrupt change from one ecosystem state to another^8^. Abrupt transitions occur when an ecosystem moves from one attraction basin into a different one in its stability landscape after surpassing a certain threshold^8^. This can have important consequences for ecosystem functioning, as the ecosystem stays in the new attraction valley – the alternative state – even if the perturbation that triggered the change has ceased. Numerous studies have demonstrated the existence of abrupt changes to alternative states in aquatic ecosystems^9,11^, terrestrial plant communities^12,13^, and the human gut microbiome in response to perturbations^14,15^. However, abrupt transitions to alterative states in soil microbial communities have so far received little attention^7^, despite their fundamental role in terrestrial ecosystems, driving key processes of organic matter decomposition, nutrient cycling, and carbon and nutrient storage^16^, which regulate ecosystem productivity. As such, abrupt transitions in soil microbial communities from one state to another may have significant implications for soil functioning, with consequences for ecosystem services, such as food production and climate regulation^1^.

Soil microbial communities are increasingly challenged by perturbations associated with human-induced environment change, including intense ‘pulse’ perturbations^17^ caused by climate extremes (e.g. droughts, heat waves and floods), which are predicted to increase in frequency and intensity as a consequence of ongoing climate change^2^. Previous studies have shown that soil microbial communities are sensitive to drought, with different microbial groups varying in their tolerance and ability to recover from drought^3,18,19^. Moreover, extreme drought has been shown to de-stabilize soil bacterial networks, potentially rendering them more vulnerable to subsequent drought^20^, although repeated droughts can also lead to soil microbial communities becoming more resistant to drought^5,21,22^. Extreme drought has been shown to shift peatland moisture characteristics^23^ and plant-soil fungi interactions^24^ to an alternative state, and repeated dry-wet cycles induced shifts in the functional state of agricultural soils, measured as soil respiration^25^.However, evidence of transitions to alternative states in soil microbial communities in response to perturbations, such as drought, is still lacking. Specifically, it remains unresolved whether increases in the intensity and frequency of drought has potential to trigger an abrupt change in soil microbial communities to an alternative taxonomic and functional microbial state.

Here, we experimentally tested whether increases in the intensity and frequency of drought triggers a shift to an alternative, functionally deleterious state in soil microbial communities. Petraitis and Dugeon^26^ proposed that experimental evidence of the existence of alternative stable states must fulfil three conditions: it must (a) demonstrate that communities differ significantly in their assembly and functional attributes within the same environment; (b) vary with the scale of the pulse perturbation, given that small perturbations are likely to be insufficient to tip the system from one state to another, whereas large perturbations are more likely to do so; and (c) be carried out over a sufficient time period to ensure that the alternative states are self-sustaining or stable. To meet these conditions, and experimentally test for the existence of alternative states in soil microbial communities, we carried out an incubation experiment in a plant-free system whereby we imposed a matrix of drought frequency and intensity treatments (one, two, three drought events and mild, intermediate and extreme drought, respectively), followed by the assessment over an extended period of time of a broad range of taxonomic and functional attributes of microbial communities in a natural grassland soil. The experimental system, although simplified, allowed us to provide experimental proof of concept of the existence of alternative states in soil microbial communities. We discovered that high intensity drought triggered a shift to an alternative microbial state of reduced structural complexity and impaired functioning. We also discovered that the functional characteristics of soil microbial communities could adapt to drought exposure, revealing a faster ability of bacterial communities to recover growth rates following drought in the alternative state.

### Changes in microbial community structure

Our findings show that extreme drought (∼11% WHC), simulating a once in a century drought in England (see methods), had a profound and long-lasting impact on soil microbial community structure and diversity, despite environmental conditions being returned to their original state (optimum moisture, 65% WHC). Six months after the end of the extreme drought, bacterial diversity (Fig 1a, b) was still reduced, while fungal Shannon diversity was significantly higher than in the non-droughted control treatment (Fid. 1d), despite no effects on fungal species richness being detected (Fig. 1c). These contrasting responses (reduced bacterial diversity but increased fungal one) are consistent with previous findings^20,27^. Bacterial and fungal community structure was strongly affected by drought (Fig. 1e, f, Extended Data Fig.1a, b) and changes in microbial community structure observed after drought were exacerbated with time. This is exemplified in the proportion of variance in community structure explained by drought intensity (Fig. 1g), which increased after 6 months compared to immediately after drought, particularly under the extreme drought treatment. However, bacterial communities in soils subjected to intermediate (23% WHC) drought, which simulated drought events in England occurring every 4 years, partially recovered (Fig. 1g), with less variance explained by drought treatments after 6 months than after drought, i.e., they were more similar to the control treatments after 6 months than at the end of the perturbation. Mild drought (40% WHC), which simulated common drought events occurring annually in England, had no detectable impact on microbial community structure. Drought frequency also significantly affected bacterial and fungal community structure (Fig. 1e, f), with those soils exposed to more frequent drought being more distinct from the non-droughted control and soil subject to fewer drought events.

**Fig. 1.**
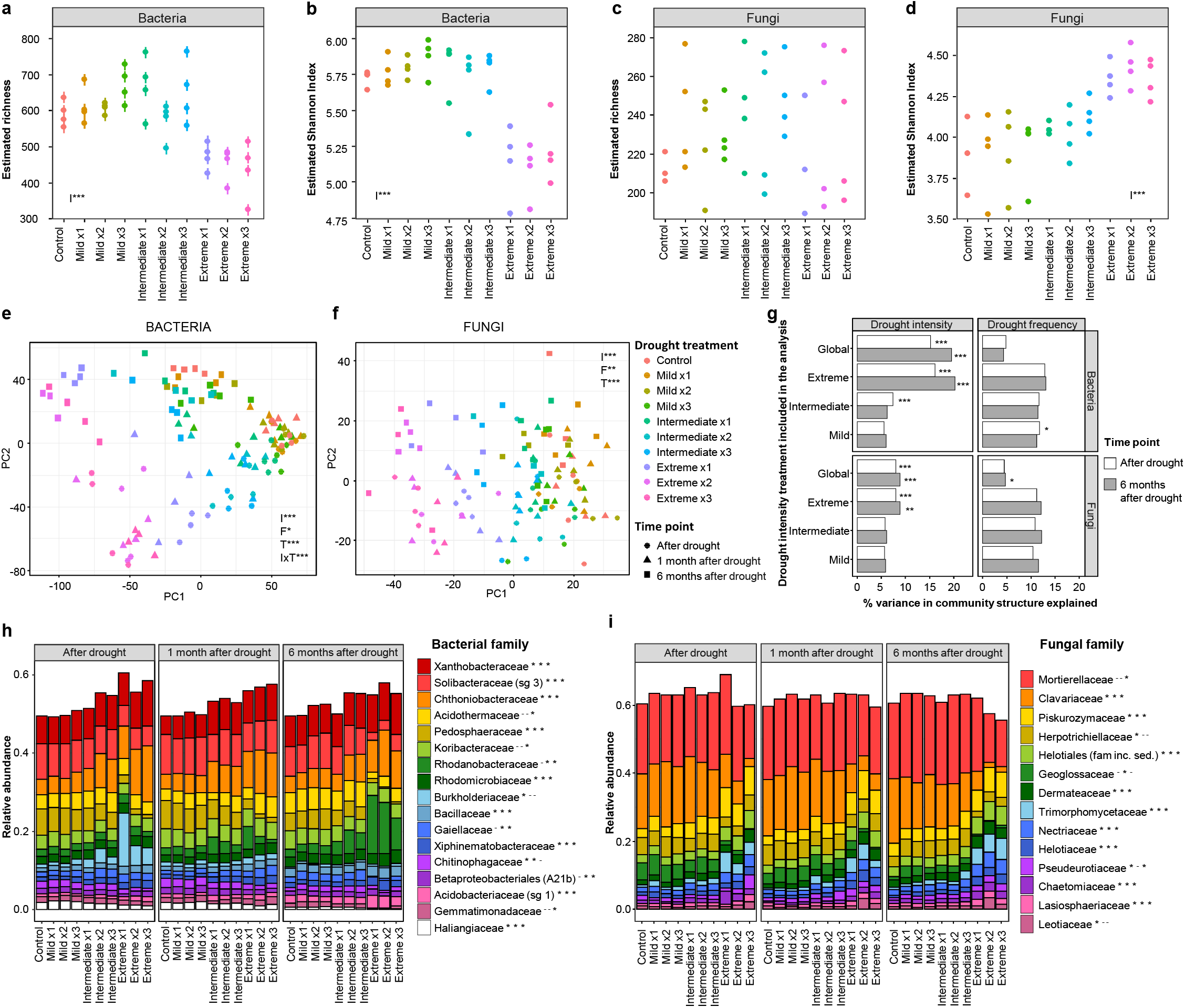
Effects of drought on soil microbial community structure. **a-d**, Alpha diversity 6 months after drought (mean ± error of estimated values). **e-f**, Community structure. The first two axes explained 17.3% (bacteria) and 5.4% (fungi) of the variance. **g**, Percentage variance explained in PERMANOVA analyses including all the treatments (global) or just one plus the control. **i-j**, Relative abundance of microbial families. Data=mean (n=4). sg = subgroup. Significance of drought intensity (I), frequency (F) and time point (T) is shown (**a-g**): *p<0.05, **p<0.01, ***p<0.001, or significance of drought treatment for each sampling time (**i-j**): ^-^ p>0.05,*p<0.05.

Sixty-eight of the 82 indicator genera identified for bacteria (Extended Data Fig. 2a) were indicators for all of the groups except the extreme drought, i.e. their abundance was significantly reduced under extreme drought. The proportion of different bacterial taxa observed in the soil communities was mainly affected by drought intensity and time, with some minor effects of drought frequency (Extended Data Table 1). Bacterial phyla Proteobacteria, Actinobacteria and Firmicutes increased in relative abundance with drought intensity, while Acidobacteria and Bacteroidetes decreased (Extended Data Fig. 1c). There was a clear shift in the bacterial community, persistent through time, from a community co-dominated by Acidobacteria and Proteobacteria in the control towards one dominated by Proteobacteria under extreme drought. Some of these Phyla-level responses were consistent with previous reports. Actinobacteria^18,21^ and Firmicutes^28,29^ have been frequently reported as drought tolerant. However, there are contrasting references to Proteobaceria, with some studies defining them as tolerant to drought^5^, and others showing a decrease in their abundance after drought^27^. On the other hand, Acidobacteria^18^ and Bacteroidetes^27^ have been described as sensitive to drought, as found in this study, although there are also some contradictory reports in the literature for both phyla^5,30^.

At family level, particularly significant were the increases in Xanthobacteraceae, Chthoniobacteraceae, Rhodanobacteraceae and a transient increase in Burkholderiaceae after drought (Fig. 1h), and the decrease in the relative abundance of Solibacteraceae and Pedosphaeraceae with drought intensity (Fig. 1h). An increase in Xanthobacteraceae in soils under drought stress has been previously reported^31^. Within the family Chthoniobacteraceae, the most abundant genus was *Candidatus Udaeobacter*. This is an extremely abundant genus in soils, adapted to grow efficiently under limited resources^32^, which could be the case in the drought exposed soils of this experiment.. The Rhodanobacteraceae family was dominated by the genus *Rhodanobacter*. This genus is common in soils and some species have been identified as plant growth promoting rhizobacteria with drought protection capacities^33^. Finally, the transient increase in Burkholderiaceae after drought was mostly driven by an increase in the genus *Massilia*. They have been identified as copiotrophic bacteria that can quickly grow under high nutrient conditions (such as those created at rewetting) but that lose their competitive advantage later on^34^, which can explain the pattern observed in this experiment. Regarding those families that decreased with drought, both agree with the literature, as Solibacteraceae^18^ and Pedosphaeraceae^35^ have been found to decrease with decreasing soil moisture.

Fungal indicator genera (Extended Data Fig. 2b) were mostly identified for the control + mild + intermediate drought group (17 out of 20), with no genus indicator for the extreme drought. The relative abundance of main fungal phyla was mostly affected by drought intensity and time (Extended Data Fig. 1e, Extended Data Table 1). We observed an increase in Ascomycota and a decrease in Mortierellomycota. In particular, we observed an increase in relative abundance of Piskurozymaceae, Helotiales, Trimorphomycetaceae and Nectriaceae, and a substantial decrease of Mortierellaceae and Clavariaceae, the two most abundant fungal families (Fig. 1i), which are both typical soil saprotrophs. In agreement with this, we observed a significant decrease with drought treatments in the relative abundance of taxa considered as saprotrophs (Extended Data Fig. 1g). Therefore, the observed increase in fungal diversity (Shannon index, Fig. 1d) was not associated with a change in species richness, but with an increase in evenness, as a result of decreasing abundances of the two major fungal taxa, Mortierellaceae and Clavariaceae. These two families have been previously identified as drought sensitive^20,36^. The observed effects could be related to the changes in nutrient availability elicited by drought (see section below). Fungi are generally considered to be more resistant to drought than bacteria^20,37^ and several studies demonstrate a lack of drought effect on fungal communities^3^. In contrast, we observed a clear change in the fungal communities, which was still evident 6 months after returning droughted soils to their original, pre-drought moisture content.

Bacterial diversity is strongly affected by soil pH at a global scale^38^ and at local scales^39^, while its impact on fungal diversity is weak or absent. In our experiment, bacterial diversity and community structure were correlated with a decrease in soil pH at the last sampling point (Extended Data Fig. 3a-c), associated with increased nitrate ion concentrations at this sampling time (Extended Data Fig. 3f). In agreement with the literature^39,40^, fungal diversity was only marginally affected by soil pH (Extended Data Fig. 3d-e).

### Changes in microbial community function

Our data show that microbial community shifts in response to extreme drought were associated with persistent changes in microbial functioning (Extended Data Fig. 3g-h). Soils exposed to extreme drought were in a significantly different functional state than the control soils even 6 months after the end of the perturbation (Fig. 2c, d).

**Fig. 2.**
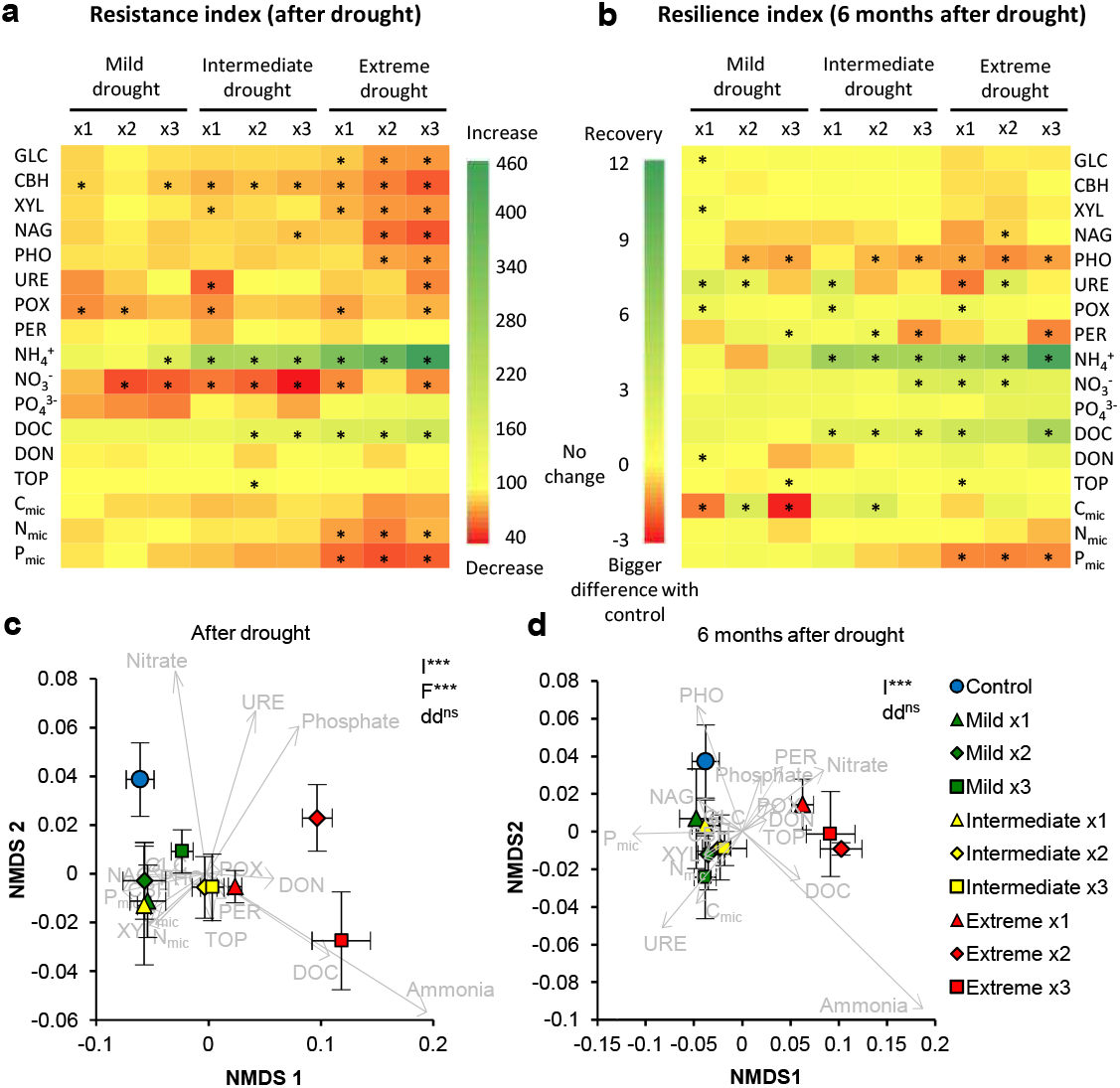
Effects of drought on soil function. **a**,**b**, Resistance and resilience indexes. Asterisks indicate significant change (value ≠ 0, p<0.05). **c**,**d**, Functional structure of soils. Mean ± error (n=4). Significance of PERMANOVA analysis evaluating the effects of drought intensity (I) and frequency (F) is shown: *p<0.05, **p<0.01, ***p<0.001. Differences in data dispersion (dd) among groups is also shown (^ns^: non-significant). GLC: *β-*glucosidase, CBH: cellobiohydrolase, XYL: xylosidase, NAG: *N*-acetylglocasiminidase, PHO: acid phosphatase, POX: phenoloxidase, PER: peroxidase, URE: urease, DOC: dissolved organic carbon, DON: dissolved organic nitrogen, TOP: total organic phosphorus, C_mic:_ microbial carbon, N_mic_: microbial nitrogen, P_mic_: microbial phosphorus.

After the end of drought, when soils were rewetted to the level of the control, seven of the eight extracellular enzyme activities evaluated were significantly reduced by drought intensity and/or frequency, displaying a low resistance to drought (Fig. 2a, Extended Data Fig. 4a-h, Extended Data Table 2). Phenoloxidase (POX) activity was slightly reduced by all drought treatments compared to the control, with no significant effect of drought intensity or frequency levels (Fig. 2a, Extended Data Fig. 4g). Only the activity of peroxidase (PER) was not affected by drought (only a marginal effect of drought frequency) (Extended Data Fig. 4h). The effects of drought intensity on soil enzymes were still detectable after six months (Extended Data Fig. 4i-p), with very little or no resilience, or even a stronger reduction of their activity than immediately after drought (phosphatase) (Fig. 2b), manifesting a persistent reduction of soil functional capacity; a functional regime shift. A reduction in soil enzymatic capacity with drought has been frequently reported^41^, probably associated with microbial death and thus reduced enzyme production, and reduced substrate diffusion that limits enzyme activity^42^. However, these effects are quite context dependent^41^ and some authors have reported increased enzyme activities under drought^43,44^, which could be related to a reduced degradation rate of soil enzymes and accumulation during dry periods^42^. In our experiment, enzyme activities correlated with available nutrients (Extended Data Fig. 3i-k). Additionally, consistently low microbial biomass after 6 months (Fig. 2b, Extended Data Fig. 3l) could explain the lack of recovery of enzymatic capacities. Ligninases (POX and PER) were less affected by drought than the hydrolytic enzymes, which could be related to the fact that fungal communities were less affected by drought than bacterial ones, as ligninases are mostly produced by fungi^42^.

Soil nutrients were highly affected by drought, particularly available ammonium and DOC, with a significant increase due to drought intensity and frequency and an interaction between them: the more intense the drought, the bigger the effect of drought frequency on their concentration (Fig. 2a, Extended Data Fig. 5a-f, Extended Data Table 2). Increases in nutrient availability after rewetting dry soils have been frequently explained by the death of microorganisms and the fact that some organic compounds in the soil become more available after drought^45,46^. On the other hand, soil nitrate concentrations were reduced with most drought treatments compared to the control, showing a low resistance (Fig. 2a). After six months, this big ammonium and DOC flush had mostly disappeared and in turn, nitrate levels significantly increased (Fig. 2b, Extended Data Fig. 5k-p). This could be related to a high nitrification activity, where the highly abundant ammonium was transformed into nitrate^47^.

Microbial biomass C, N and P were reduced by increasing drought intensity, and this effect was still detected after 6 months of returning soils to moisture levels of the control, with microbial P in extreme drought treatments showing a stronger decrease with time (Fig 2b, Extended Data Fig. 5g-i, q-s, Extended Data Table 2). Decreases in microbial biomass due to drought have been widely reported^27,48^, although some authors observed an increase in microbial biomass under drought^49^. The constant decline of microbial P over time could be related to reduced phosphatase activity (Extended Data Fig. 3m). This is in agreement with the study of Dijkstra and collaborators^50^ that showed a strong reduction in microbial P uptake under drought. However, a decrease in microbial biomass over time in bare soils is expected, as there is no additional C input from plants into the system.

### Adaptation of microbial growth characteristics to drought

Our assessment of the responses of microbial communities with a history of drought to a further drying/rewetting cycle helped us examine legacy effects of drought and adaptation of growth responses, as a community, of bacteria and fungi. Drought intensity had a legacy effect resulting in shorter lag times and higher cumulative growth (Fig. 3a, b) after rewetting, indicating a faster recolonization ability. The increased and faster bacterial growth in soils with a drought history has been previously observed^22^, and can be a good adaptive strategy in soils exposed to frequent drought events. On the other hand, cumulative fungal growth (Fig. 3c) significantly decreased with previous drought intensity, which could underpin the reduced abundance of the two dominant fungal families. This result contrasts with other studies where fungal growth was not affected by drought history^19,51^. However, the fungal growth capacity in our study could have also been constrained by the high bacterial growth in the soils^52^. These effects on microbial growth were not dependent on the microbial biomass of soils before drying/rewetting (Extended Data Fig. 3n,o). Cumulative respiration after rewetting significantly decreased with previous drought intensity (Fig. 3d), and thus it seems to be driven by fungi, as it follows approximately the same pattern as cumulative fungal growth. This contrasts with some recent observations, where respiration was mostly driven by bacterial growth^19^. Alternatively, this decrease in cumulative respiration with previous drought intensity could be also related to resource availability. Previous extreme drought led to a strong increase of DOC, which was mostly used one month after drought, and this likely depleted the soil carbon available to fuel a respiration peak after the additional drying/rewetting cycle. Reduced carbon availability in soils after drought has been previously reported^47^, as well as less intense respiration peaks after repeated drying/rewetting cycles^22,47^.

**Fig. 3.**
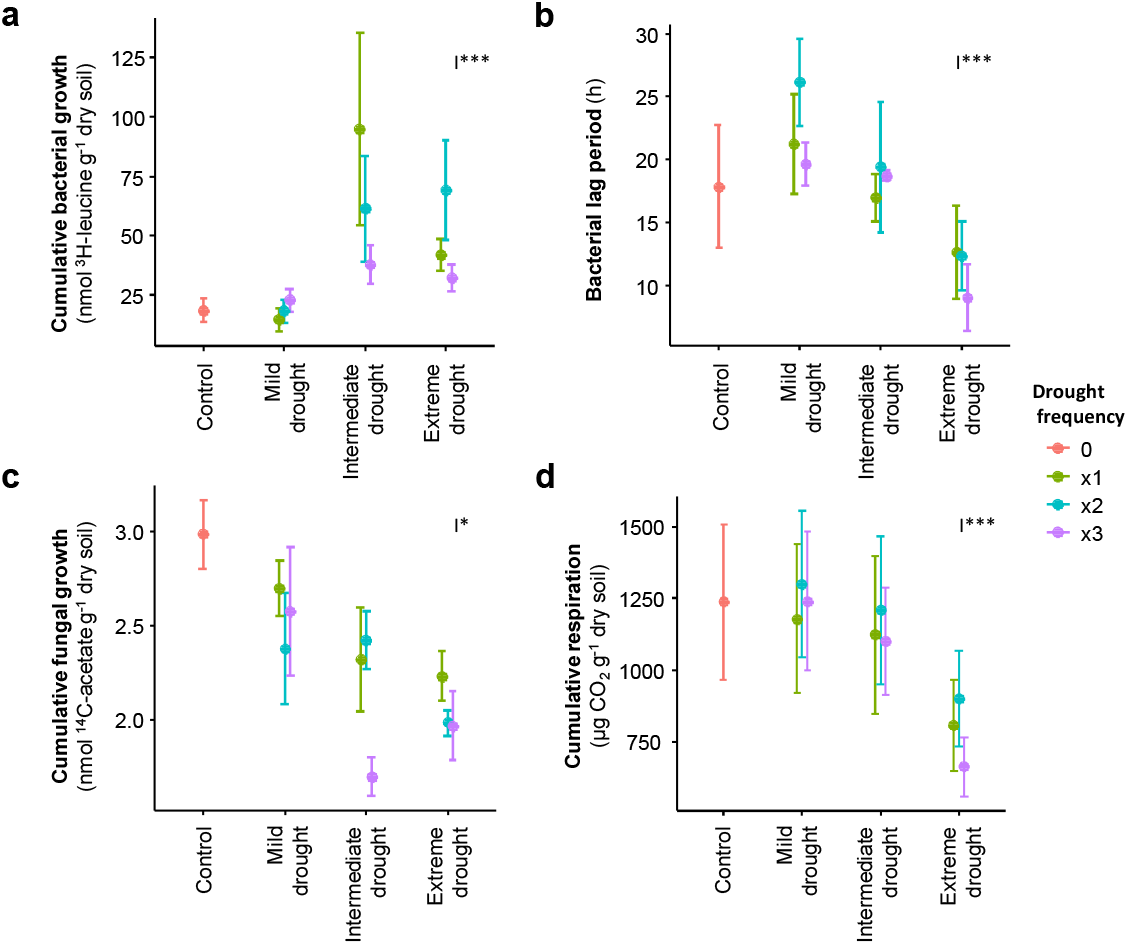
Adaptation to drought of microbial community traits. Growth responses to a further drying/rewetting cycle of soils with different drought history, one month after drought. Significance of LME evaluating the effects of drought intensity (I) and frequency (F) is shown: *p<0.05, **p<0.01, ***p<0.001.

Fungi were more resistant to low moisture than bacteria, as shown by the IC10 (moisture level at which growth rate is reduced by 10%), but this was not significantly affected by previous drought treatments (Extended Data Fig. 6). However, there were some trends evident, with bacteria and fungi responding in opposite directions. While previous exposure to drought had a tendency to reduce bacterial resistance to drought (higher IC values), fungi tended to show a higher resistance to drought under some of the previous treatments (lower IC values), although this was quite variable. Respiration showed reduced resistance to drought with previous drought intensity.

We observed legacy effects of drought on soil microbial communities, as previously described. For example, pre-exposure to drought has been demonstrated to increase bacterial resistance^5,21^ and resilience^22^ to subsequent droughts, although others have reported the opposite pattern, with previous drought damaging microbial communities in the long-term^20,27^. We observed an increased recolonization capacity of bacteria, albeit a lower tolerance to reduced soil moisture, while fungi showed a potential increased resistance to drought but with less cumulative growth upon rewetting. Thus, there seems to be a trade-off between growth after rewetting, which can be interpreted as resilience, and resistance to low moisture, as previously proposed^53^. These strategy changes could be the result of a shift in the relative abundance of different taxa within the community or due to changes in individual taxa’s physiology or traits (evolution). In any case, the response of fast re-coloniser bacterial taxa appears to shape bacterial communities after drought, as they occupy empty niches left after drought, conditioning community assembly afterwards^54^.

### Drought threshold and shift to alternative state

Our findings provide experimental evidence that the harshness of drying determined the response pattern of microbes upon rewetting. Although gradient designs are best suited to evaluate thresholds^55^, the design of the present experiment with four levels of soil moisture provides evidence of a potential threshold for a shift to an alternative microbial state at a water content of ∼15% WHC (∼9% volumetric soil water content), which corresponds to a 30 year recurrent drought event in England (Extended Data Fig. 6d-g). Below this moisture level, soil microbial communities were considerably and persistently restructured with impaired functioning, and they failed to recover over a period of 6 months, despite returning moisture levels to those of the control. The existence of this threshold is further supported by a previous study that demonstrated that below a threshold of 14% WHC, the growth pattern of bacteria upon rewetting changed significantly^56^.

To further evaluate potential shifts to alternative states in our experiment, we mapped microbial community composition and function with multivariate analysis, as it is a useful tool to visualise stability landscapes and regime shifts^57^. We observed a clear pattern in both community structure and function of soil microbial communities where soils subjected to extreme drought occupy a clearly distinct space separated from the control, which can be interpreted as an alternative basin in the stability landscape, an alternative state of the system (Extended Data Fig. 6h-k). Moreover, our experiment met most of the recognised criteria for detecting an alternative state^26^. First, we demonstrated the existence of two different microbial communities given the same environmental condition (optimum soil moisture during the period after drought). Second, different intensities of drought (different scale of pulse perturbation) were found to have contrasting effects on soil microbial communities, with mild and intermediate drought showing reversible effects, but persistent changes in response to extreme drought. Third, our experiment was conducted over an extended period of time, representing the typical length of the growing season in the north of England, where the soil samples were collected. The identified microbial shifts are unlikely to represent a “stable” alternative state. Nevertheless, we experimentally demonstrate that microbial communities subject to extreme drought shift towards a different state that is distinct from its original one and also from the non-droughted control soils. Therefore, we propose that the detected microbial shifts reflect an alternative “transient” state^54^ or simply an alternative state.

## Conclusion

Our findings experimentally demonstrate that extreme and frequent drought triggers a shift to an alternative microbial state, with pronounced and persistent reduction in soil microbial community functional capacity, modified taxonomical composition of reduced complexity, and faster ability of bacterial communities to recover growth rates following drought. Moreover, we show that this state transition was driven by drought intensity and occurred at a threshold of ∼15% WHC, although more frequent and extreme events produced stronger effects than milder and less frequent drought events. Future studies are needed to consider the role of extrinsic factors that might modify the vulnerability of soil microbial communities to drought-induced transitions to alternative states, such as the presence of plants, which can help the system recover after drought^58^, or differences in nutrient availability and other soil abiotic properties^7^. Nevertheless, our results provide experimental evidence for the existence of alternative taxonomic and functional states in soil microbial communities driven by extreme drought, with potentially deleterious consequences for soil health.

## METHODS

### Soil collection and experimental design

A pot experiment was designed to test the effects of drought intensity and frequency (3 levels each, full-factorial including a well-watered control), on the microbial communities in grassland soils. Soils were collected in Selside, Yorkshire Dales (54.17 N, 2.34 W), from four independent plots (replicates). Turf was removed (2-3 cm), and topsoil collected from 3-10 cm depth. Main soil characteristics are shown in Extended Data Table 3. Soils were sieved (4 mm mesh) and divided into pots (8 cm height, 8 cm diameter, and 170 ml volume), filled to mimic the mean field bulk density (0.82 g cm^-3^). Pots were incubated at 18 ºC, 30% air relative humidity, and kept at 65% water holding capacity (WHC), which correspond to ∼40% volumetric water content. After 3 weeks of stabilisation, drought treatments were applied by different watering regimes: control treatments received 100% of the water loss, mild drought treatments received 2/3, intermediate drought received 1/3 and extreme drought received no water. Pots were watered every other day with autoclaved water. Soil moisture was evaluated gravimetrically. Drought lasted 2 weeks followed by 2 weeks of recovery when pots were slowly rewetted to optimum moisture (65% WHC). Drought was repeated up to 3 times depending on the drought frequency treatment. Control pots were always kept at 65% WHC. Drought levels were compared for contextualisation to historical data of soil moisture (1991-2020) in England, made available by the CCI SM project^59,60^. Minimum annual values of soil moisture were fitted to an extreme value distribution (Gumbel distribution) with the package “extRemes”^61^. The maximum drought experienced by treatments in each cycle was approximately 40% WHC for mild drought (∼23% volumetric water content), which corresponded to the average values of soil moisture during the summer (Jun-Aug), 23% WHC for intermediate drought (∼14% volumetric water content), which resembled common summer drought events (once every 4 years), and 11% WHC for extreme drought (∼7% volumetric water content), which simulated a once in a century drought. Extended Data Fig. 7a summarizes the evolution of WHC. WHC capacity was affected by the drought treatments (Extended Data Fig. 7b), so after each drought cycle, WHC was re-evaluated. The decrease of moisture retention capacity of soils after repeated drought has previously been documented^23,62^.

Immediately after the last drought cycle, samples were collected to evaluate the resistance of the system. We consider the perturbation the entire drying/rewetting cycle, and thus, to evaluate the resistance we harvested the pots once the perturbation had ended (when all pots recovered the same WHC, 11 days after the start of the rewetting, Extended Data Fig. 7a). Additionally, pots were harvested over time, to evaluate the resilience of the system in the long term (1, 3 and 6 months after drought), which covers the length of the typical growing season for the study sites where soil was collected in the Yorkshire Dales (Met Office UK). 40 extra pots were harvested 1 month after drought to evaluate the adaptation of microbial growth characteristics to drought. Total number of pots: 200.

### Microbial community structure

Soil samples were collected in Eppendorf tubes (approx. 0.25 g) and frozen at -80 ºC just after sampling. DNA was extracted in frozen samples, without thawing, with PowerSoil DNA isolation Kit (Qiagen, Germany). DNA was sent to Macrogen sequencing service (Macrogen Inc., Korea), for Illumina sequencing (MiSeq v3). Fungal diversity was evaluated by ITS2 sequencing, using the primer pair 5.8S-Fun and ITS4-Fun^63^. Bacterial diversity was evaluated by 16S V3-V4 sequencing, with primers Bakt_341F and Bakt_805R^64^. Microbial community analysis was not done in samples from the 3 months after drought time point. Alongside the samples, three extraction blanks were included and a mock community sample for each primer pair: 19 strains genomic DNA even mix from Bakker lab for fungi^65^, and MSA-1000™ 10 strain even mix genomic material for bacteria from the American Type Culture Collection (ATCC, Manassas, US).

The quality of sequences was assessed using FASTQC^66^. Primers were removed using cutadapt^67^ (fungi) or obitools^68^ (bacteria). Low quality areas at the end of the reads were trimmed using truncLen= c(275,200) (fungi) or truncLen= c(260,180) plus 10 additional nucleotides at the beginning of forward reads (bacteria). Sequences were analysed using the DADA2 pipeline^69^ with default parameters, except maxEE for fungi which was set to 4 in each direction. After filtering and de-noising steps 77% (fungi) or 44% (bacteria) of the reads were retained, and 4,227 (fungi) or 16,924 (bacteria) amplicon sequence variants (ASV) were identified. Taxonomic identification was performed by IDTAXA taxonomic classification method in DECIPHER^70^ package using UNITE reference database (sh_general_release_dynamic_s_01.12.2017.fasta) for fungi and SILVA database (SILVA_SSU_r132_March2018.RData) for bacteria. Database was refined by four consecutive steps. 1) Renormalization to counteract the problem of tag-switching^71^: for the abundant ASVs (≥ 100 reads), eliminate the reads of the samples corresponding to a cumulative frequency of less than 3% for each particular ASV. 2) Lulu algorithm to reduce the number of erroneous ASVs and achieve more realistic biodiversity metrics^72^, setting the minimum ratio at 2 for fungi and minimum match at 99% for bacteria, and the rest parameters set as default. The election of these parameters was based on the results of the mock community sample. 3) Minimal abundance filtering, by removing any reads that represent < 0.02% abundance in each sample. 4) Blank correction, where ASVs were removed if the total abundance in blanks divided by the total abundance in sample was greater than 10%, in each particular ASV. Finally, only fungi or bacteria sequences were retained. For fungal database, other eukaryotic reads that were removed represented the 0.01% of the reads. For bacteria, taxa unclassified at domain level represented the 2.69% of the reads, Archaea reads represented the 0.004% and chloroplast sequences represented the 0.0007%. All of them were removed. Final databases contained 2,760 ASVs and 4,316,693 reads for fungi, and 7,198 ASVs and 2,392,480 reads for bacteria. Sampling depth varied from 25,873– 53,007 (mean 36,275) reads in fungi and 12,595–58,150 (mean 19,937) reads in bacteria, and it was equally distributed among drought treatments. Rarefaction curves were plotted using the *rarecurve* function in “vegan”^73^, and these indicated that sampling depth was sufficient for all samples, plateauing at around 7,500 reads.

### Soil enzymes

*β-*glucosidase (GLC), cellobiohydrolase (CBH), xylosidase (XYL), *N*-acetylglocasiminidase (NAG) and acid phosphatase (PHO) were measured photometrically according to Jackson *et al*. 2013^74^, with modifications. 3.75 g of sieved soil was suspended in 5 mL of sodium acetate buffer (50 mM, pH 5.0). 150 μl of soil slurry were introduced into a 96-well deepwell block and mixed with 150 μl of a saturating substrate solution: 25 mM *p*NP-*β-*glucopyranoside for GLC, 2 mM *p*NP-*β-*D-cellobioside for CBH, 25 mM *p*NP-*β-*xylopyranoside for XYL, 5 mM *p*NP-N-acetyl-*β-*D-glucosaminide for NAG and 5 mM *p*NP-phosphate disodium salt hexahydrate for PHO. Plates were incubated at 18 ºC for 0.5 h (PHO), 1.5 h (GLC), 3.5 h (XYL and NAG) or 4 h (CBH) under continuous shaking. Blocks were centrifuged (2900 *g*, 5 min), 100 μl of the supernatant pipetted into transparent 96-well plates and mixed with 200 μl of 50 mM NaOH solution. Absorbance was measured at 405 nm in a plate reader (EZ400 Research, Biochrom, Germany) and corrected for soil and substrate colouration. Reported activity is the mean of four analytical replicates.

Urease (URE) was measured by the optimised high throughput method^75^ without modifications. Absorbance was evaluated in a microplate reader (EZ400 Research, Biochrom, Germany) and reported activity is the mean of four analytical replicates. Phenoloxidase (POX) and peroxidase (PER) activities were measured photometrically following Sinsabaugh & Linkins 1988^76^ method, with modifications. 0.25 g of soil were suspended in 25 ml of sodium acetate buffer (50 mM, pH 5.0). 0.4 ml of soil slurry were extracted under continuous shaking and mixed (1:1) with a 20 mM L-3,4-dihydroxyphenylalanin (L-DOPA) solution in a deep-well block. Blocks were shaken for 10 min and centrifuged (2900 *g*, 5 min). 250 μl of the supernatant were pipetted into transparent 96-well plates. For peroxidase activity, wells additionally received 10 μl of a 0.3% H_2_O_2_ solution. Absorbance was measured at 450 nm in a plate reader (EZ400 Research, Biochrom, Germany) at the starting time point (t_0_), after 1.5 h (t1 for PER) and after 20 h (t1 for POX) at 18 ºC. Enzyme activity was calculated from the difference in absorption between the two time-points divided by L-DOPA molar extinction coefficient (7.9 μmol^-1 77^). For PER activity, POX activity was subtracted. Reported activity is the mean of three analytical replicates.

### Nutrient pools

Different nutrient pools were measured by means of soil extractions with different extracting solutions depending on the nutrient and the pool of interest^78^. All soil nutrients were evaluated in duplicates, and reported values are the mean of those two analytical replicates. Dissolved organic carbon (DOC) and dissolved organic nitrogen (DON) were evaluated in water extracts (5 g soil in 28 ml MiliQ water) and plant available nitrogen (ammonium and nitrate) were evaluated in KCl extracts (2.5 g soil in 12.5 ml 1M KCl). In both cases, soil with extracting solutions were horizontally shaken at 200 rpm for 30 minutes and filtered through Whatman nº 42 filter papers (KCl extracts) or 0.45 μm syringe filters (H2O extracts). N pools were measured in AA3 HR Auto Analyser (Seal Analytical, UK) while C pools were measured in 5000A TOC-L analyser (Shimadzu, Japan). Plant available P was extracted with acetic acid solution (1 g soil + 25 ml 2.5% acetic acid) and detected by molybdate colorimetry using AA3 HR Auto Analyzer (Seal Analytical, UK).

Organic P was estimated by evaluation of available phosphate before and after sample ignition^79^. Two portions of dry soil (2 g) were weighed. One of them was burned at 550ºC for 4 h in a furnace. Afterwards, both samples were extracted with 50 ml 0.5 M H2SO4 by horizontally shaking at 120 rpm for 16 h. Extracts were filtered (Whatman No. 42) and phosphate was evaluated by the ascorbic acid method^79^. After sample neutralization with NaOH, the colour reaction was carried out and the amount of blue was evaluated by absorbance at 880 nm in a CLARIOstar plate reader (BMG Labtech, Germany).

Microbial biomass C and N was measured using the fumigation–extraction techniques^80,81^. 2.5 g of soil were fumigated with CHCl3 for 24 h. Soluble C and N were extracted from the fumigated and from un-fumigated samples with 12.5 ml 0.5 M K2SO4. Soil + extracting solution were shaken and filtered (Whatman No. 42) and total C and N were analysed in TOC and AA, respectively. Microbial C and N flush (difference between fumigated and un-fumigated samples) were converted to microbial biomass using kEC factor of 0.35^82^ and kEN factor of 0.54^80^.

Microbial P was estimated with the hexanol fumigation and extraction with anion-exchange membranes method^83^. Membrane strips (1 × 4 cm) were prepared by shaking in 0.5 M NaHCO3. For each sample, three portions of fresh soil (2 g) were weighed into 50 ml tubes: unfumigated, fumigated and spiked samples. In each tube, 30 ml deionized water and three anion-exchange membrane strips were added. Fumigated samples received 1 ml of hexanol while spiked samples received 1 ml of a 20 mg ml^-1^ P solution. All tubes were shaken for 24 h. The membranes were then removed and rinsed with deionized water and phosphate recovered by shaking for 1 h in 20 ml of 0.25 M H2SO4, with detection at 880 nm by automated molybdate colorimetry using AA3 HR Auto Analyzer (Seal Analytical, UK). Microbial phosphorus were calculated as the difference between the fumigated and un-fumigated samples, corrected by the sorption percentage (spiked samples) and transformed using a kEP factor of 0.40^84^.

### Adaptation of microbial growth characteristics to drought

Adaptation of soil bacterial and fungal community characteristics to drought was evaluated by assessing their growth and respiration rate responses to a subsequent drying/rewetting cycle in soil samples that had been recovering for one month after drought. We did two complementary tests: first, we assessed the response to a full drying/rewetting cycle, to compare the ability of microbial communities to recover after rewetting and secondly, we evaluated moisture dependences to test the ability of microbes to withstand a lack of moisture. For the first test, soil samples were completely air dried for two days and then rewetted to 60% WHC. After rewetting, microbial growth rates were measured with a high temporal resolution of approximately 6 h for a 160 h period (more frequent at the beginning and every 24 h afterwards, 12 time points in total). Bacterial and fungal growth rates were assessed via radioisotope incorporation^85^. Bacterial growth rates were evaluated by ^3^H-leucine incorporation into proteins^86^ while fungal growth rates were measured by ^14^C-acetate incorporation into ergosterol^87^. To measure soil respiration, 1.0 g of soil was weighed into 20-ml glass vials, which were purged with pressurised air, sealed with crimp caps and incubated for 6 h in the dark at 17°C. CO2 production was measured using a gas chromatograph equipped with a methaniser and an FID detector. To evaluate the moisture dependence of microbial processes, microbial growth and respiration rates were measured during the drying down of the soils. Then, IC10 (i.e. the moisture level at which growth and respiration rates were reduced by 10%) was determined, which allowed comparison of the drought tolerance of the different microbial processes^22^.

### Statistical analyses

All statistical analyses were done in R v4.0^88^. To evaluate microbial community structure, we investigated alpha diversity, community structure with ordination analyses, proportion of different taxa and functional guilds (only for fungi) and we performed indicator species analysis. Due to the issues associated with using rarefied or relative abundance data for diversity tests and differential abundance analyses^89^, different approaches or data transformations were applied for each analysis.

Alpha diversity was calculated estimating the unobserved diversity^90^. Richness was estimated with “breakaway” package^91^ and Shannon index with “DivNet” package^92^. Community structure was explored with a variance stabilization transformation (VST) of data, using “DeSeq2” package^93^. PERMANOVA analyses using Euclidean distance of transformed data were performed to evaluate differences among drought treatments, with soil as random factor, using *adonis* function in “vegan” package^73^. To better visualise the effects of the drought treatments, a correction to eliminate the soil effect was performed with *removeBatchEffect* function in “limma” package^94^. A PCA analysis was then performed in the corrected database using “vegan” ^73^. To predict functional guilds and trophic modes from the fungal taxonomic data we used FUNGuild^95^. Only those assignments considered highly probable or probable were used, while possible assignments were discarded. Assignments with multiple and contradictory guilds to the same taxa were also discarded. In average, 29.1% of the taxa (774 ASVs) were assigned and used in this analysis. Of those, 82 were assigned to order level, 158 to family level and 534 to genus level. For relative abundance graphs, only taxa (phylum, order or family) with relative abundances >1% across all samples were plotted.

Indicator species analysis was performed using *multipatt* function in the “indicspecies” package^96^ to identify bacterial and fungal taxa associated with drought and control treatments. We tested for indicator taxa depending on the intensity of drought, comparing the control treatment with the extreme drought and some specific combinations (control+mild, control+mild+intermediate, and intermediate+extreme drought). Prior to analysis, ASVs were agglomerated at genus level using *tax_glom* in “phyloseq” package^97^, and libraries were normalised using cumulative sum scaling (CSS)^98^, using *phyloseq_transform_css* function in “metagMisc” package^99^. CSS-normalised abundance heatmaps were produced using *plot_heatmap* function in “phyloseq” package^97^.

Resistance and resilience of soil functions were evaluated with a linear regression analysis between the value of the variable under drought and time after drought (in weeks). We used relative data in percentage (i.e., value drought pot/value control pot × 100), and we performed individual analyses for each of the nine drought intensity and drought frequency combinations. The value of the intercept was used as a resistance index (RS), as it represents the relative difference of the drought treatment with the control just after drought. RS can be directly interpreted as the % change compared to the control treatment. Additionally, we used the value of the slope as a resilience index (RL). This value was adjusted depending on the direction of change after drought with the formula: if intercept < 100, then RL = slope; if intercept > 100, then RL = −slope. Thus, a positive RL indicates a recovery of the system; variables in drought pots are getting more similar to those in control pots. On the other hand, RL = 0 reflects no change (no resilience), with drought pots showing the same difference with the control ones that what it was at the end of the drought. Moreover, a negative RL can be interpreted as delayed response to drought or a continuous negative effect even when the perturbation has finished, as negative RL mean a bigger difference with the control over time than what it was at the end of the drought. RL values can be directly interpreted as percentage recovery/change per week.

Soil functional data (soil extracellular enzymes and nutrient pools) were used in a non-metric multidimensional scaling (NMDS) ordination analysis, performed with the function *metaMDS* in “vegan”^73^, using relative data (value/average of the control at each sampling date). Multifunctionality index^100^ was calculated with all the soil enzymatic activities, which represent the organic matter decomposition capacity of soils. Data were standardised by z transformation separately for each sampling time, which removed overall differences between harvest time points^101^. Subsequently, the average of all standardised values was used as the multifunctionality index.

The effects of drought intensity and frequency in all variables were analysed by linear mixed effect models (LME) with drought intensity and frequency as fixed factors and soil as random factor, with *lme* function in “nmle” package^102^. To obtain a balanced design, control pots were removed from the analysis. Normality of the residuals and homoscedasticity were confirmed with Anderson-Darling test and Levene test, respectively. If needed, natural logarithmic or square root transformations were applied. If the assumptions were still not met, Kruskal-Wallis test was performed instead. *P* values after multiple comparisons (for bacterial and fungal taxa) were adjusted with the Benjamini-Hochberg adjustment^103,104^ using *p*.*adjust* function.

## ACKNOWLEDGMENTS

This work was supported by a grant from the Ramon Areces Foundation and UK Biotechnology and Biological Sciences Research Council (BBSRC) Discovery Fellowship (BB/S010661/1) awarded to IC, and a BBSRC grant (BB/I009000/2) awarded to RDB. The work was also supported by a grant from the Knut and Alice Wallenberg foundation (KAW 2017.0171) awarded to JR. We are grateful to members of the Soil and Ecosystems Ecology Laboratory who helped with the running of the experiment, especially Debbie Ashworth and Diego Abad Martín, and Eva Berglund at Lund University. We also thank Lokeshwaran Manoharan, Marina Semchenko, Océane Nicolitch, Hayley Craig, Arthur Broadbent and Chris Sweeney for help with data analyses, and Dr. Matthew Bakker for proving the fungal mock community used for ITS2 sequencing.

## AUTHOR CONTRIBUTIONS

IC and RDB conceived the study and designed the experiment, with inputs from JR. IC performed nutrient analyses, enzyme assays, and microbial sequencing work. AL and LH performed microbial growth rates measurements. IC analysed the data and wrote the manuscript, with significant contributions from all authors.

