## Supplementary Material for "Extreme drought triggers transition to an alternative soil microbial state"

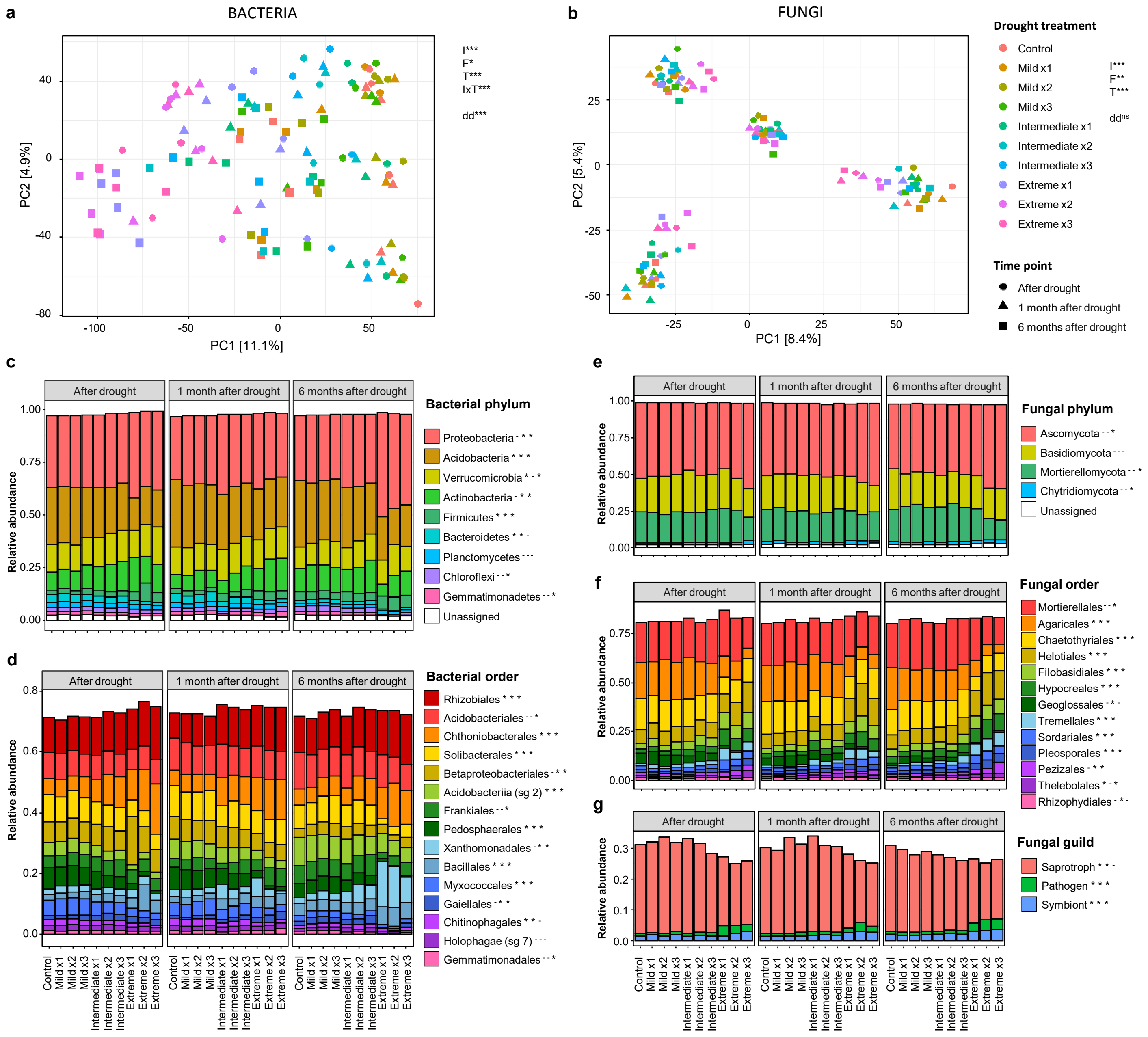


**Extended Data Fig. 1 | Microbial community structure.** Data based on 16S (bacteria) and ITS2 (fungi) amplicon sequencing an summarised by drought intensity and frequency (x1: 1 event, x2: 2 events, x3: 3 events) treatments. **a,b,** Community structure evaluated by principal component analysis of VST transformed data, without removing soil effect. Significance of PERMANOVA analysis evaluating the effects of drought intensity (I), frequency (F) and sampling time (T), with soil as random factor, is shown in each graph: *p<0.05, **p<0.01, ***p<0.001. Differences in data dispersion (dd) among groups is also shown ($^{ns}$: non-significant). Variance explained by soil is 9.9% for bacteria (**a**) and 18.4% for fungi (**b**). **c-g**, Relative abundance of different microbial taxa depending on drought treatment and sampling time. Data = mean of 4 replicates. Only taxa with relative abundances >1% across all samples are plotted. Fungal guild data extracted from FUNGuild database. Significance of linear mixed models evaluating the effects of drought treatment for each sampling time with soil as random factor, is shown to the right of each taxa name as - p>0.05,*p<0.05. sg = subgroup.

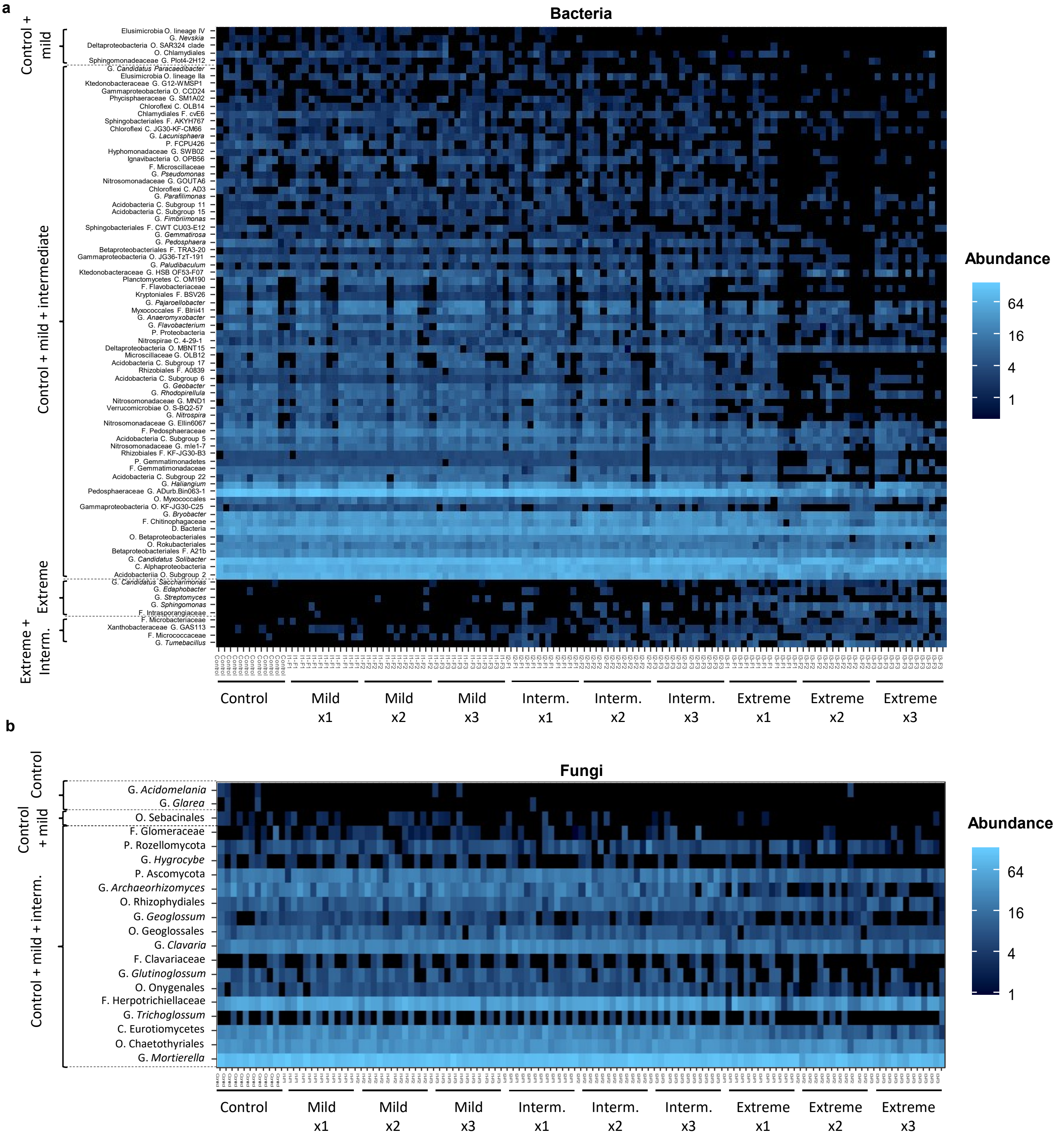


**Extended Data Fig. 2 | Indicator species analysis.** Heatmaps showing cumulative sum scaling (CSS) normalised abundances of soil bacteria (**a**) and fungi (**b**) indicator taxa identified for different drought intensity and frequency (x1: 1 event, x2: 2 events, x3: 3 events) treatments. Only taxa significant at $p \leq 0.001$ (bacteria) or $p \leq 0.01$ (fungi) are shown. Data agglomerated at genus level for analysis. ASVs unassigned at genus level, show the name of the lowest assigned taxonomical rank (P=Phylum, C=Class, O=Order, F=Family, G=Genus). Interm. = intermediate.

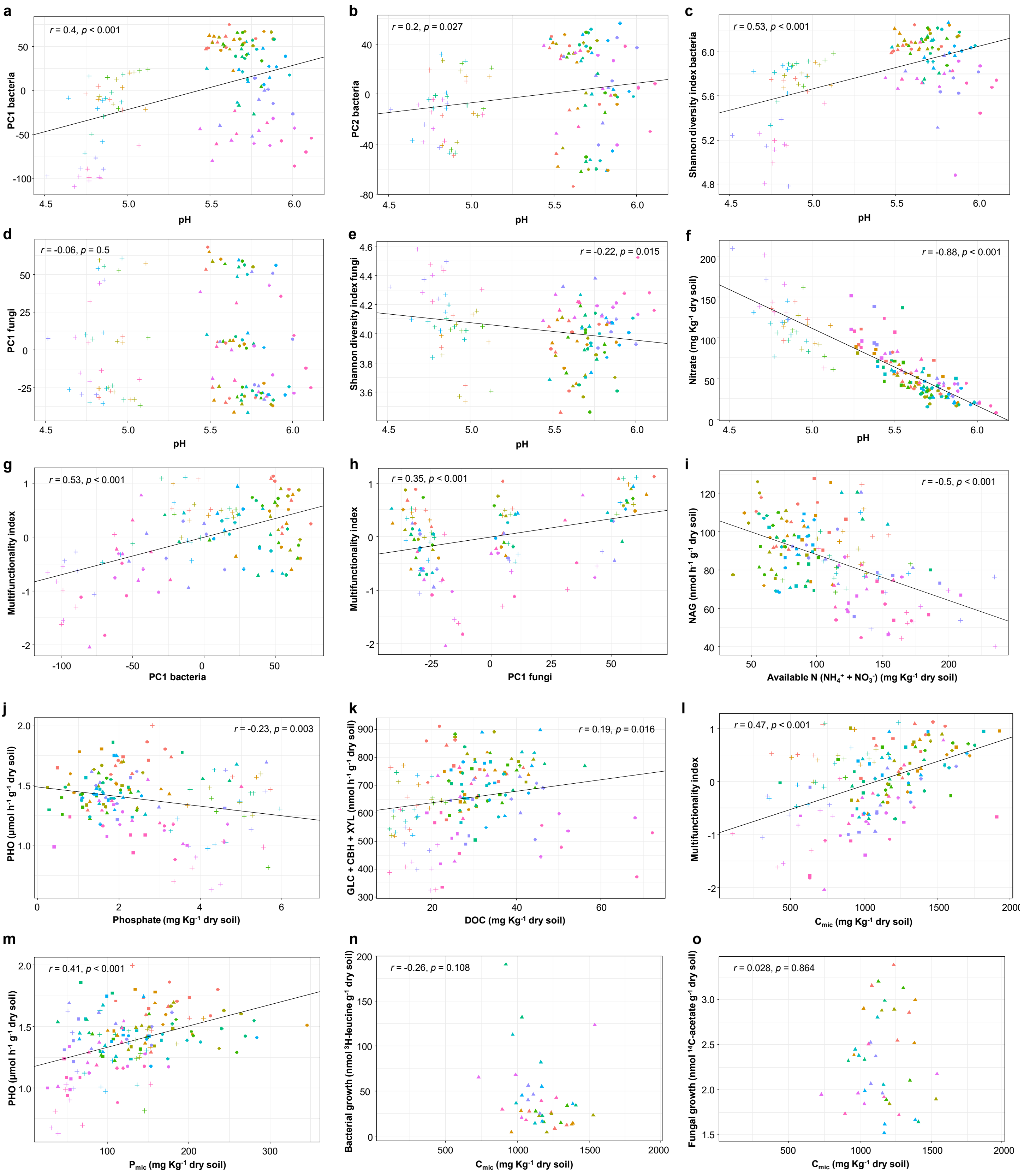


**Extended Data Fig. 3 | Correlations between soil chemical and biological variables.** Pearson correlation value and significance test are shown in each graph. Regression line is represented when the correlation is significant ($p < 0.05$). GLC: *β*-glucosidase, CBH: cellobiohydrolase, XYL: xylosidase, NAG: *N*-acetylglocasiminidase, PHO: acid phosphatase, $C_{mic}$: microbial carbon, $N_{mic}$: microbial nitrogen, $P_{mic}$: microbial phosphorus. PCA axis correspond to the microbial community structure analyses in Extended Data Fig. 1. Multifunctionality index represents a combined index of all analysed enzyme activities.

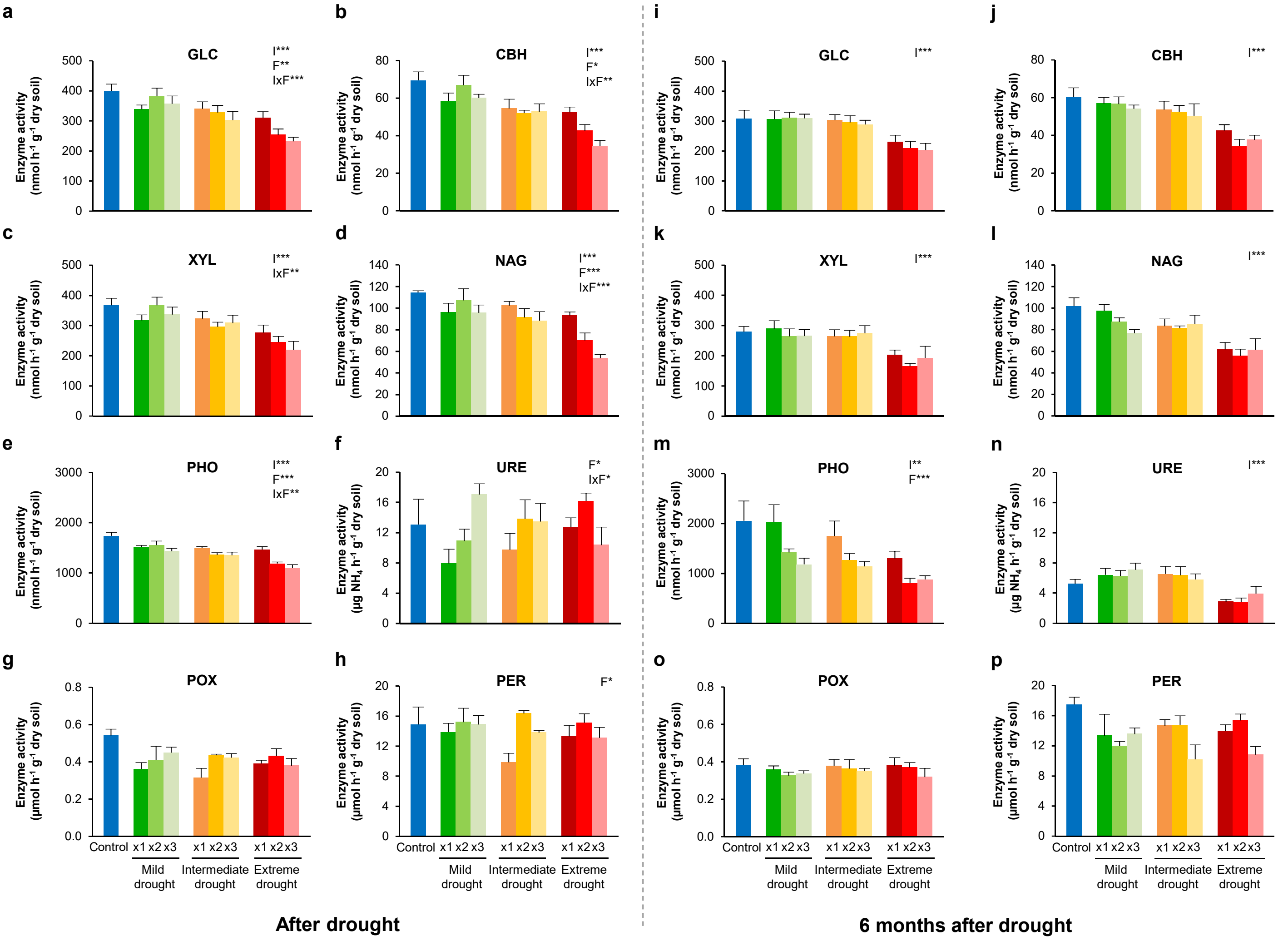


**Extended Data Fig. 4 | Effects of drought on soil enzyme activities.** Data after drought (**a-h**) and after 6 months afterwards (**i-p**), summarised by drought intensity and frequency (x1: 1 event, x2: 2 events, x3: 3 events) treatments. Significance of linear mixed models evaluating the effects of drought intensity (I) and frequency (F), with soil as random factor, is shown in each graph: *p<0.05, **p<0.01, ***p<0.001. Values = mean ± standard error, n=4. GLC: *β*-glucosidase, CBH: cellobiohydrolase, XYL: xylosidase, NAG: *N*-acetylglocasiminidase, PHO: acid phosphatase, URE: urease, POX: phenoloxidase, PER: peroxidase.

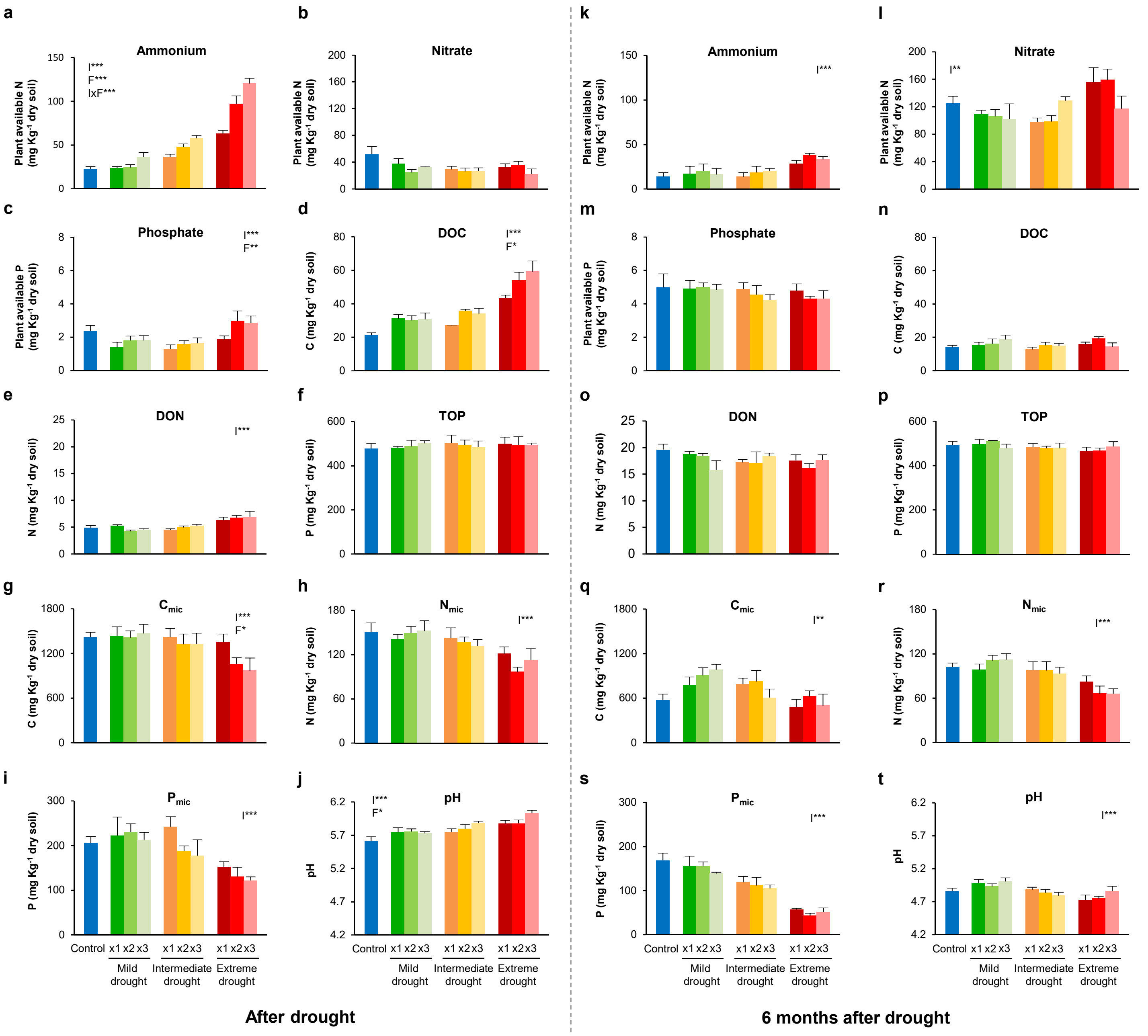


**Extended Data Fig. 5 | Effects of drought on soil nutrients, microbial biomass and pH.** Data after drought (**a-j**) and after 6 months afterwards (**k-t**), summarised by drought intensity and frequency (x1: 1 event, x2: 2 events, x3: 3 events) treatments. Significance of linear mixed models evaluating the effects of drought intensity (I) and frequency (F), with soil as random factor, is shown in each graph: *p<0.05, **p<0.01, ***p<0.001. Values = mean ± standard error, n= 4. DOC: dissolved organic carbon, DON: dissolved organic nitrogen, TOP: total organic phosphorous, $C_{mic}$: microbial carbon, $N_{mic}$: microbial nitrogen, $P_{mic}$: microbial phosphorus.

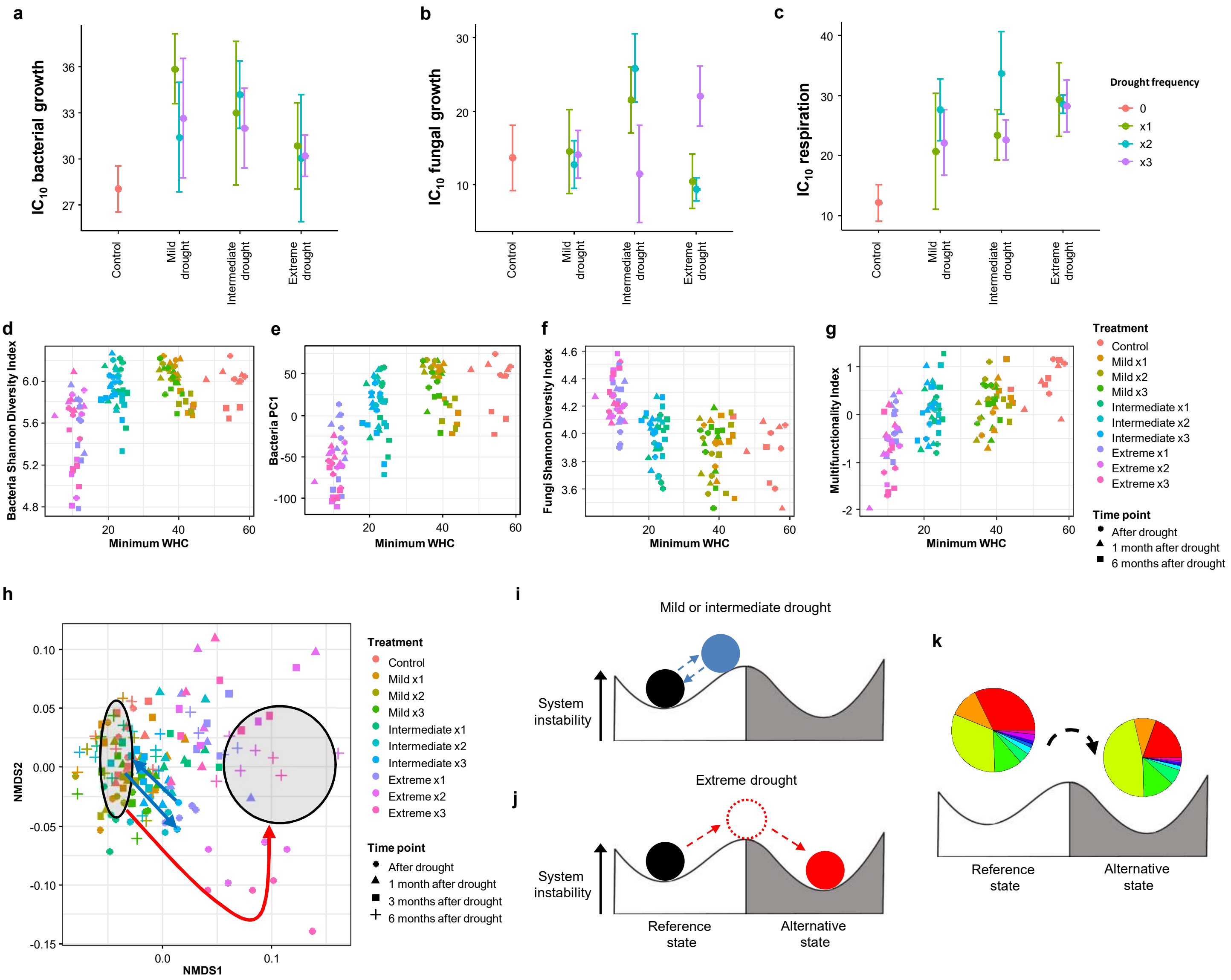


**Extended Data Fig. 6 | Drought legacy effects, drought thresholds and shift to an alternative state in soil microbial communities. a-c** Moisture dependence of bacterial and fungal growth and respiration depending on drought history. $IC_{10}$ represents the moisture level (in % water holding capacity) at which growth and respiration rates are reduced 10%. Drought intensity and drought frequency had no significant effects on these variables. Drought intensity 1=mild, 2=intermediate, 3=extreme; drought frequency 1=1 event, 2=2 events, 3=3 events. **d-g,** Drought threshold: microbial community diversity, composition and multifunctionality depending on minimum water holding capacity (WHC) experienced by soils. **e**, Functional capacity of soils evaluated by principal non-metric multidimensional scaling (NMDS) analysis with schematic representation of a shift to an alternative state. Mild or intermediate drought (blue arrows) moved the system from the reference state (circle on the left), but they bounced back. Extreme drought (red arrow) moved the system further away, crossing a threshold, into an alternative state (circle on the right). **f-h**, Diagrammatic representations of the effects of drought on the system, represented as a ball moving up and out of the stability basin. Mild or intermediate drought do not push the ball far enough, bouncing back to the reference state (**f**), while extreme drought pushes the ball out of the reference state into a different stability basin, an alternative state (**g**), with distinct functionality and microbial community composition (**h**).

a

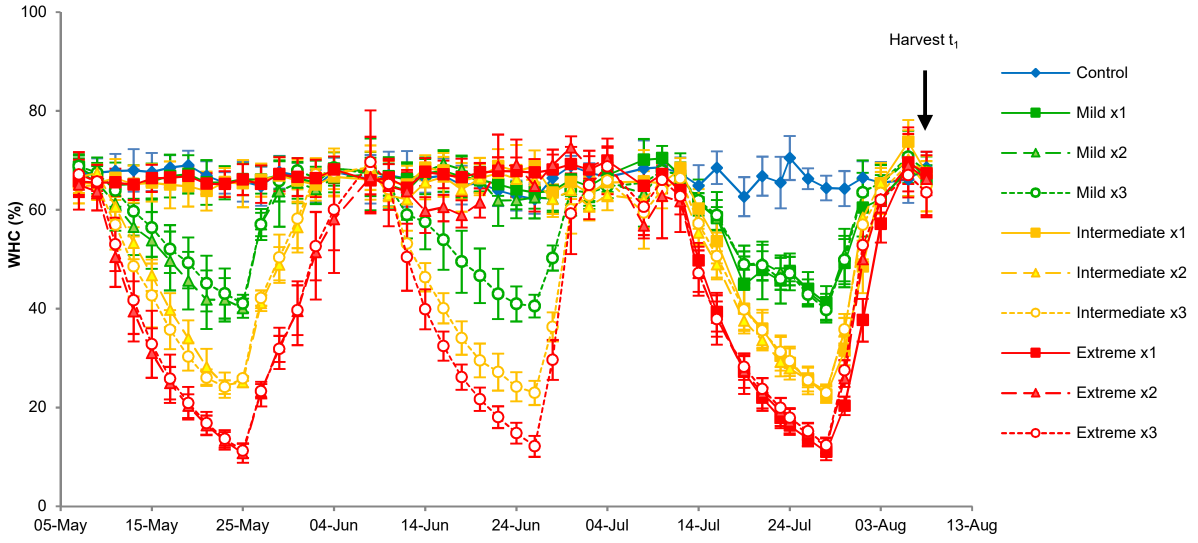


b

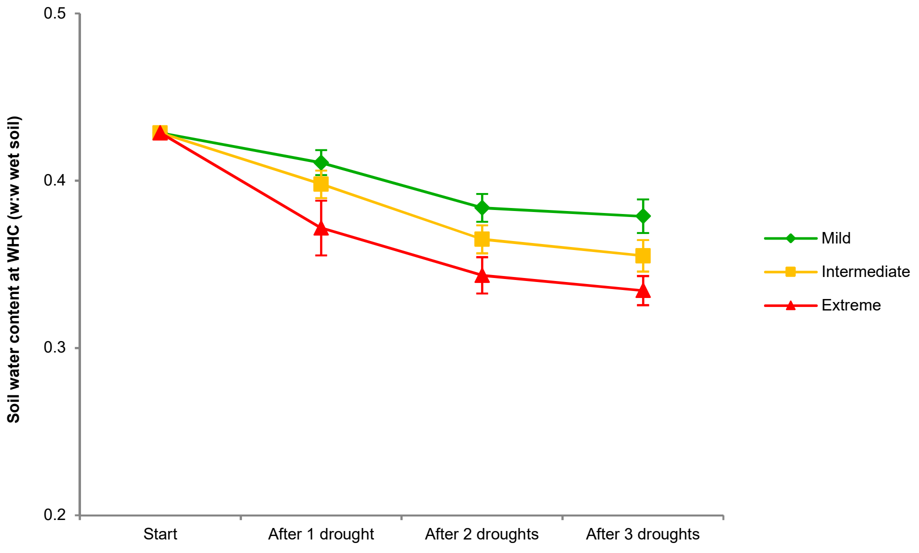


**Extended Data Fig. 7 | Soil water holding capacity (WHC). a,** Evolution of WHC along the experimental period depending on drought intensity (I) and frequency (F). The arrow indicates the date of the harvest after drought. **b,** Effects of drought on WHC of soils. Values = mean ± standard deviation, n= 4.

**Extended Data Table 1**. Effects of drought intensity, drought frequency, and harvest time point on soil bacterial and fungal taxa, analysed by linear mixed models with soil as random factor. F values and significance levels are given as *<0.05; **<0.01; ***<0.001; $^{ns}$: not significant.

| | Intensity (I) | | Frequency (F) | | Time point (T) | | I x F | | I x T | | F x T | | I x F x T | |
|---|---|---|---|---|---|---|---|---|---|---|---|---|---|---|
| *Bacterial Phylum* | | | | | | | | | | | | | | |
| Proteobacteria | 22.23 | *** | 4.50 | * | 7.49 | ** | 1.53 | ns | 14.28 | *** | 0.38 | ns | 0.18 | ns |
| Acidobacteria | 116.89 | *** | 1.85 | ns | 35.68 | *** | 0.93 | ns | 5.34 | *** | 1.18 | ns | 1.63 | ns |
| Verrucomicrobia | 13.02 | *** | 5.60 | ** | 19.26 | *** | 1.82 | ns | 0.94 | ns | 3.68 | ** | 1.54 | ns |
| Actinobacteria | 3.83 | * | 4.01 | * | 2.44 | ns | 0.63 | ns | 6.78 | *** | 1.15 | ns | 0.89 | ns |
| Firmicutes | 11.64 | *** | 0.47 | ns | 4.13 | * | 1.21 | ns | 1.08 | ns | 0.76 | ns | 0.57 | ns |
| Bacteroidetes | 66.46 | *** | 1.47 | ns | 65.46 | *** | 0.83 | ns | 5.10 | ** | 1.21 | ns | 1.04 | ns |
| Planctomycetes | 11.10 | *** | 0.49 | ns | 159.63 | *** | 0.72 | ns | 0.45 | ns | 0.81 | ns | 1.66 | ns |
| Chloroflexi | 9.31 | *** | 1.61 | ns | 1.05 | ns | 0.43 | ns | 9.24 | *** | 1.53 | ns | 0.56 | ns |
| Gemmatimonadetes | 8.50 | *** | 1.86 | ns | 10.72 | *** | 0.38 | ns | 9.86 | *** | 1.07 | ns | 0.45 | ns |
| *Fungal Phylum* | | | | | | | | | | | | | | |
| Ascomycota | 16.46 | *** | 6.24 | ns | 0.07 | ns | 5.73 | *** | 1.72 | ns | 0.36 | ns | 0.75 | ns |
| Basidiomycota | 4.27 | * | 2.06 | ns | 3.46 | * | 1.63 | ns | 0.52 | ns | 1.22 | ns | 0.24 | ns |
| Mortierellomycota | 9.77 | *** | 1.41 | ns | 1.44 | ns | 3.03 | * | 5.27 | *** | 1.14 | ns | 1.03 | ns |
| Chytridiomycota | 8.18 | *** | 0.30 | ns | 5.56 | ** | 0.36 | ns | 0.10 | ns | 3.05 | * | 1.21 | ns |

**Extended Data Table 2**. Effects of drought on soil extracellular enzyme activities and soil nutrients, analysed by linear mixed models with drought intensity (I) and frequency (F) as fixed factors and soil as random factor. It is also shown if transformation to meet model assumptions was needed (Ln = natural logarithm transformation, Sqrt= square root transformation). Significant results are highlighted in bold. GLC: *β*-glucosidase, CBH: cellobiohydrolase, XYL: xylosidase, NAG: *N*-acetylglocasiminidase, PHO: acid phosphatase, URE: urease, POX: phenoloxidase, PER: peroxidase, DOC: dissolved organic carbon, DON: dissolved organic nitrogen, TOP: total organic phosphorous, $C_{mic}$: microbial carbon, $N_{mic}$: microbial nitrogen, $P_{mic}$: microbial phosphorus.

| | **After drought (t1)** | | | | | | | **Six months after drought (t4)** | | | | | | |
|---|---|---|---|---|---|---|---|---|---|---|---|---|---|---|
| **Variable** | **Intensity (I)** | | **Frequency (F)** | | **IxF** | | | **Intensity (I)** | | **Frequency (F)** | | **IxF** | | |
| | F | p | F | p | F | p | | F | p | F | p | F | p | |
| GLC | 67.58 | **<0.001** | 8.62 | **0.002** | 8.19 | **<0.001** | | 50.44 | **<0.001** | 0.83 | 0.449 | 0.44 | 0.779 | |
| CBH | 44.29 | **<0.001** | 5.07 | **0.015** | 6.18 | **0.002** | | 49.93 | **<0.001** | 2.31 | 0.121 | 0.95 | 0.455 | |
| XYL | 49.03 | **<0.001** | 1.94 | 0.166 | 5.01 | **0.004** | | 21.47 | **<0.001** | 1.06 | 0.363 | 0.43 | 0.787 | |
| NAG | 34.43 | **<0.001** | 13.60 | **<0.001** | 6.85 | **<0.001** | | 19.13 | **<0.001** | 1.12 | 0.342 | 1.18 | 0.344 | |
| PHO | 32.69 | **<0.001** | 20.20 | **<0.001** | 5.70 | **0.002** | | 8.03 | **0.002** | 11.43 | **<0.001** | 0.43 | 0.782 | |
| URE | 0.28 | 0.757 | 3.51 | **0.046** | 3.31 | **0.027** | | 35.91 | **<0.001** | 0.61 | 0.554 | 0.94 | 0.451 | |
| POX | 0.14 | 0.872 | 2.84 | 0.078 | 0.88 | 0.493 | | 0.54 | 0.589 | 1.59 | 0.226 | 0.35 | 0.840 | |
| PER | 0.90 | 0.419 | 5.34 | **0.012** | 1.61 | 0.203 | | 0.13 | 0.877 | 2.85 | 0.078 | 1.81 | 0.160 | |
| $NH_4^+$ | 165.39 | **<0.001** | 33.77 | **<0.001** | 7.39 | **<0.001** | | 23.35 | **<0.001** | 2.51 | 0.102 | 0.58 | 0.683 | |
| $NO_3^-$ | 1.00 | 0.382 | 2.18 | 0.135 | 2.40 | 0.078 | | 7.45 | **0.003** | 0.14 | 0.871 | 2.33 | 0.085 | |
| $PO_4^{3-}$ | 17.09 | **<0.001** | 6.12 | **0.007** | 0.91 | 0.472 | | 1.36 | 0.274 | 0.98 | 0.391 | 0.28 | 0.889 | |
| DOC | 38.76 | **<0.001** | 3.81 | **0.037** | 1.56 | 0.219 | Ln | 1.32 | 0.285 | 1.03 | 0.371 | 0.94 | 0.459 | |
| DON | 16.68 | **<0.001** | 0.14 | 0.867 | 1.55 | 0.219 | Ln | 0.17 | 0.848 | 0.27 | 0.765 | 1.37 | 0.274 | |
| TOP | 0.06 | 0.945 | 0.02 | 0.983 | 0.27 | 0.892 | | 2.58 | 0.096 | 0.13 | 0.882 | 1.33 | 0.286 | |
| $C_{mic}$ | 15.66 | **<0.001** | 4.02 | **0.031** | 2.38 | 0.080 | | 7.87 | **0.002** | 0.77 | 0.474 | 0.88 | 0.493 | |
| $N_{mic}$ | 22.02 | **<0.001** | 0.77 | 0.474 | 1.79 | 0.164 | | 12.34 | **<0.001** | 0.06 | 0.944 | 0.93 | 0.464 | |
| $P_{mic}$ | 15.86 | **<0.001** | 2.38 | 0.114 | 0.75 | 0.570 | | 42.82 | **<0.001** | 0.43 | 0.654 | 1.18 | 0.343 | Sqrt |
| pH | 20.42 | **<0.001** | 5.43 | **0.011** | 2.20 | 0.099 | | 11.10 | **<0.001** | 0.63 | 0.542 | 1.30 | 0.297 | |

**Extended Data Table 3**. Main soil chemical characteristics. Variables measured after pot experiment setup but before drought treatments. Mean ± standard deviation are shown (n=4). OM: organic matter content. DOC: dissolved organic carbon, DON: dissolved organic nitrogen, TOP: total organic phosphorus, $C_{mic}$: microbial carbon, $N_{mic}$: microbial nitrogen, $P_{mic}$: microbial phosphorus.

| **Variable** | **Value** |
|---|---|
| pH | 6.11 ± 0.08 |
| OM (%) | 8.89 ± 0.07 |
| Plant available $NH_4^+$ (mg $Kg^{-1}$ dry soil) | 8.09 ± 1.28 |
| Plant available $NO_3^-$ (mg $Kg^{-1}$ dry soil) | 6.20 ± 1.40 |
| Plant available $PO_4^{3-}$ (mg $Kg^{-1}$ dry soil) | 1.83 ± 0.64 |
| DOC (mg $Kg^{-1}$ dry soil) | 49.28 ± 4.45 |
| DON (mg $Kg^{-1}$ dry soil) | 5.52 ± 1.73 |
| TOP (mg $Kg^{-1}$ dry soil) | 556.37 ± 58.29 |
| $C_{mic}$ (mg $Kg^{-1}$ dry soil) | 1348.4 ± 259.4 |
| $N_{mic}$ (mg $Kg^{-1}$ dry soil) | 172.0 ± 35.9 |
| $P_{mic}$ (mg $Kg^{-1}$ dry soil) | 182.5 ± 34.7 |
